# raxtax: A k-mer-based non-Bayesian Taxonomic Classifier

**DOI:** 10.1101/2025.03.11.642618

**Authors:** Noah A. Wahl, Georgios Koutsovoulos, Ben Bettisworth, Alexandros Stamatakis

**Affiliations:** Biodiversity Computing Group, Institute of Computer Science, Foundation for Research and Technology Hellas, 100 Nikolaou Plastira, 70013 Heraklion, Crete, Greece; Computational Molecular Evolution Group, Heidelberg Institute for Theoretical Studies, Schloss-Wolfsbrunnenweg 35, 69118 Heidelberg, Baden-Württemberg, Germany; Institute for Theoretical Informatics, Karlsruhe Institute of Technology, Kaiserstraße 12, 76131 Karlsruhe, Baden-Württemberg, Germany

**Keywords:** taxonomic classification, barcoding, biodiversity, sequence analysis, evolutionary biology

## Abstract

**Motivation:** Taxonomic classification in biodiversity studies is the process of assigning the anonymous sequences of a marker gene (barcode) or whole genomes (metagenomics) to a specific lineage using a reference database that contains named sequences in a known taxonomy. This classification is important for assessing the diversity of biological systems. Taxonomic classification faces two main challenges: first, accuracy is critical as errors can propagate to downstream analysis results; and second, the classification time requirements can limit study size and study design, in particular when considering the constantly growing reference databases. To address these two challenges, we introduce `raxtax`, an efficient, novel taxonomic classification tool for barcodes that uses common *k*-mers between all pairs of query and reference sequences. We also introduce two novel uncertainty scores which take into account the fundamental biases of reference databases.

**Results:** We validate `raxtax` on three widely used empirical reference databases and show that it is 2.7-100 times faster than competing state-of-the-art tools on the largest database while being equally accurate. In particular, `raxtax` exhibits increasing speedups with growing query and reference sequence numbers compared to existing tools (for 100,000 and 1,000,000 query and reference sequences overall, it is 1.3 and 2.9 times faster, respectively), and therefore alleviates the taxonomic classification scalability challenge.

**Availability and Implementation:** `raxtax` is available at https://github.com/noahares/raxtax under a CCNC-BY-SA license. The scripts and summary metrics used in our analyses are available at https://github.com/noahares/raxtax_paper_scripts. The source code, sequence data and summarized results of the analyses are available at https://doi.org/10.5281/zenodo.15057027.

## 1 Introduction

Biodiversity researchers frequently need to address the question: Which species are present in my sample? A common solution consists in identifying and subsequently sequencing a well-conserved region of the genome which is present in all organisms under study (1–3). Such regions, known as barcodes (4), are then used to identify species. The ribosomal 16S gene, the cytochrome oxidase 1 (COX1), and the internal transcribed spacer (ITS) regions are examples of frequently used barcodes in distinct regions of the tree of life (see, e.g., (5–7)). As using barcodes for DNA-based species identification constitutes a routine analysis task, there exist several widely used taxonomic classification tools, such as SINTAX (8), IDTAXA (9), the RDP Naive Bayesian classifier (RDP) (10), and BayesANT (11). These highly cited tools, deploy distinct algorithmic approaches to determine the species that are present in a sample.

The major design and one major quality criterions for any taxonomic classification tool are: assign sequences quickly and correctly. Species identification accuracy is critical, as it typically constitutes the first step in biodiversity analyses. Therefore, errors are likely to be propagated to downstream analyses and results. However, we are in the midst of the next generation sequencing data avalanche which is being further intensified by an increasing number of biodiversity field studies (12, 13). The amount of data being generated has outpaced Moore’s law for the last decade (14). Hence, we need to perform barcoding sequence data analysis more efficiently. Otherwise, biodiversity research will be increasingly constrained by the computational resources available.

To alleviate this scalability challenge we introduce a novel tool, which we call `raxtax`, and demonstrate that it is at least as accurate as the widely-used existing tools SINTAX, IDTAXA, RDP, and BayesANT. Furthermore, we demonstrate that `raxtax` is 2.7 to 100 faster in comparison to the competing tools listed.

`raxtax` achieves high accuracy in conjunction with computational efficiency via a *k*-mer based matching approach. That is, we formulate sequence similarity as follows: Compute the expected number of matching *k*mers between the reference sequence and a random sampling of the *k*-mers of a query sequence. The key insight is that if a query sequence is more similar to a reference sequence, the number of expected matching *k*-mers will be higher. Other tools have used analogous sampling techniques to great effect (e.g., MetaCache in the context of metagenomic studies (15, 16)). Here, instead of sampling k-mers, we devise an analytical solution. With this reformulation of the problem we can derive closed analytical solutions that allow for computing the exact probability that a given reference sequence is (among) the best matches for a random sample of query sequence *k*-mers. Given a set of DNA reference sequences (each with a taxonomic annotation), `raxtax` computes the best-match probabilities for each anonymous query sequence, and reports the best matching lineages with their per-rank confidence scores by aggregating these probabilities at each taxonomic rank (clade). Finally, we also use these per-rank confidence scores to compute uncertainty scores for each assignment of a query to a lineage. Each of these quantities and their interpretations are discussed in Section 2. `raxtax` is available as open source code and pre-compiled binaries at https://github.com/noahares/raxtax under a CC-NC-BY-SA license.

## 2 Method

Given a sequence *𝒮* (consisting of characters from the set *{*`A`, `C`, `G`, `T`*}*), a *k*-mer is a sub-sequence *𝒮* [*i*..*i* + *k*], *i ∈* [|*𝒮*| *− k*] of length *k*. The set of *k*mers, *Q*, associated with includes all unique *k*-mers of *𝒮*. For our current implementation of `raxtax`, we fix *k*:= 8 to allow for some computational optimizations (see Section 3.1), but in principle the method can be adapted to any *k*.

Strictly matching *all k*-mers of each query sequence against *all* reference sequences is not only time and memory intensive, but also highly sensitive to sequencing errors (17). On the other hand, only matching a small random sample of *k*-mers does not constitute an appropriate solution either. In particular, if the reference sequences are highly similar and/or share a large fraction of *k*-mers, numerous repetitions with small random samples will be required to distinguish between plausible assignments and therefore increase run-times. Instead, we use a combinatorial approach for selecting a random subset of *k*-mers from the query to match against the reference. This allows to obtain accurate results while being computationally efficient at the same time.

Assume that we are given the set of all *k*-mers *Q* which have been extracted from a query sequence and that we intend to match them against a set of reference sequences *D* = *{D*_1_, …, *D*_*n*_*}*. For each *D*_*i*_ there exists a corresponding set of all *k*-mers contained therein, denoted by *𝒦*_*i*_. Let *𝒦* = {*𝒦*_1_, …, *𝒦*_*n*_*}* be the set of all *k*-mer sets. To find the best matching *𝒦*_*i*_ for a given *Q*, we need to identify the *K*_*i*_ which maximizes the expected number of matches from a random sampling of *t k*-mers from *Q*. We label this sample as *S*_*t*_(*Q*). Define *P*_*i*_ as the probability that the reference *k*-mer set *𝒦*_i_ has the most *k*-mers in common with a random sampling of *Q*, or more formally:

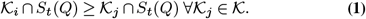

Our method for computing this probability is described in Sections 3.3 and 3.4.

Define the probability that a reference *k*-mer set *𝒦*_*i*_ has *m* matching *k*-mers with *S*_*t*_(*Q*) as

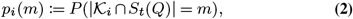

*p*_*i*_(*m*):= *P* (|*K*_*i*_ *∩ S*_*t*_(*Q*)| = *m*), **(2)** which is a probability mass function (PMF). Using this definition, we can now compute the cumulative mass function (CMF) by marginalizing over the possible match sizes that are indexed by *l*. Then, we take the product over the other references indexed by *j* to compute the probability of no other reference having more than *m* matches,

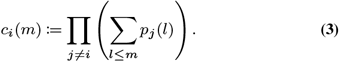

The probability that *𝒦*_*i*_ is among the best matches, given a sample size *t* then is

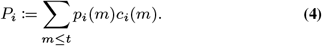

Additionally, we normalize the values in *P* via the *L*1 norm in order to compute *confidence (scores)*. This operation simplifies the subsequent confidence accumulation at different taxonomic ranks. As a result, the reported values are not, strictly speaking, probabilities. Instead, they report the confidence regarding the relative ranking of reference for matching a query.

Given a clade *B* of the reference taxonomy, we define the confidence of *B* being among the best matches relative to other clades of the same rank as

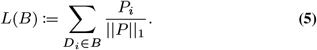

To simplify the notation, we define *ℒ* (*D*_*i*_) as the lineage confidence vector for reference sequence *D*_*i*_. *ℒ* (*D*_*i*_) is a sequence of *L*(*·*) values for the taxonomic lineage where *𝒜*_*i*_ is a series of nested partitions (clades) of the reference sequences (*D*_*i*_ = *𝒜*_0_ *⊆* … *⊆𝒜*_*i*_ *⊆* … *⊆ D*). An example lineage tree with a lineage confidence vector for a reference sequence *D*_4_ is shown in Figure 1.

**Fig. 1.**
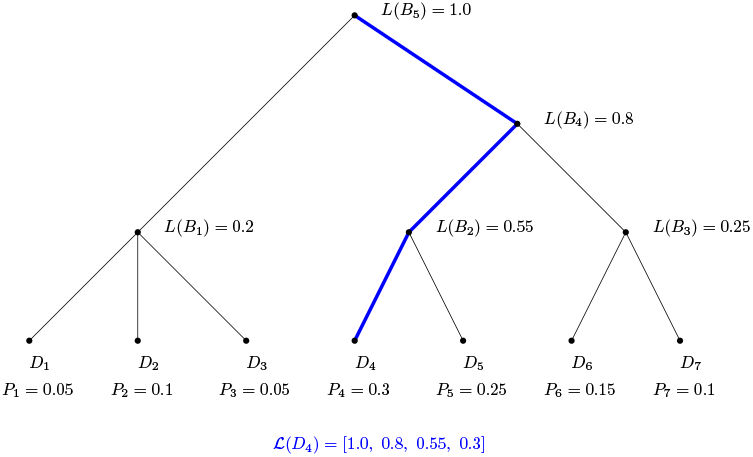
A simple lineage showing how a *ℒ*-vector is constructed. The contributors to *ℒ* (*D*_4_) are highlighted in blue.

### 2.1. Uncertainty scores

The per-rank confidence values *L*(*·*) that we compute with `raxtax` will be biased by the taxonomic distribution of reference sequences in the database. Because the values at high-level ranks are the sum over all per-sequence values within those ranks, interpreting a confidence value of 0.5 requires knowledge about the relative frequency of that clade in the reference database. For instance, consider the case that one family represents 50% of the database. In this case, by chance alone, a substantial proportion of the total confidence score will be assigned to reference sequences in this over-represented family. Therefore, to better interpret the confidence values relative to the reference database properties, we report two additional uncertainty scores.

Let 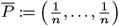 be the *expected* confidence vector for a sequence that is highly dissimilar (i.e., *k*-mer set intersections will be of approximately the same size) to all reference sequences. In analogy to using *L* for *P* values (Equation 5), we define *L* as the *expected* confidence of obtaining a higher-level rank assignment based on *P*. This means that the expected values of higher-level ranks represent the potential database bias. We will use these values to derive an uncertainty score for the global (*per-sequence*) and local (*per-rank*) assignment signals, i.e., the deviation of the observed confidence values from the expected values based on the reference database bias.

#### The *local assignment signal*

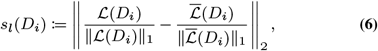

quantifies the uncertainty in *ℒ* (*D*_*i*_) as the Euclidean distance between the computed and expected per-rank confidence values (with normalization). Analogously, we define the *global assignment signal*

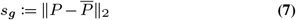

to quantify the reference sequence level confidence scores as the Euclidean distance between the computed and expected per-sequence confidence values. We describe how to interpret and use the local and global assignment signals in Supplement 6.5.

## 3 Implementation

`raxtax` is written in Rust (compiled with version 1.76) and is parallelized over the query sequences using the `rayon` library (18). In this section we describe the algorithmic techniques and data structures we use to optimize`raxtax.`

### 3.1. Calculating Intersection Sizes

To compute the match scores for all query-reference pairs, we need to compute the intersection of the two *k*mer sets. Because computing intersection sizes accounts for at least half of the processing time of a query it is important to optimize them. A naïve implementation requires computing *𝒪* (*nm*) intersections, where *n* is the number of query sequences, and *m* is the number of reference sequences.

The best case run time for a sorted set intersection of sets *A* and *B* is *O*(*min*(|*A*|, |*B*|)) via a linear scan when *A ⊆ B*.

While there exist numerous fast set intersection algorithms (19), most pairs of *k*-mer sets satisfy |*A ∩ B*| *≪ min*(|*A*|, |*B*|). Hence, it will be more efficient to ask which reference sequences contain a specific *k*-mer and store these results in a lookup table. This lookup table is computed once for all *k*-mers and reference sequences and is query-independent. It can therefore be saved for any analyses that use the same reference database. Given this lookup table, we simply perform a lookup of the *k*-mers in the query sequence to compute the intersection of a query-reference pair. Thereby, we reduce the work for one query-reference pair from *O*(*min*(|*A*|, |*B*|)) to *O*(|*A ∩ B*|) where *A* and *B* are the respective *k*-mer sets.

Because we discard *k*-mers that include gaps and ambiguous characters, they can be represented in a memory-efficient manner by only using two bits per DNA character. By setting *k*:= 8, we can thus uniquely store an 8-mer in a 16-bit unsigned integer (`u16`) by using its corresponding bit representation. While parsing the reference sequences, we create a lookup table that for each 8-mer (represented as a `u16`) holds a sorted list of reference sequences that contain it. When extracting the *k*-mers from a query sequence later-on, we can use this lookup table to rapidly identify those reference sequences that contain each query sequence *k*-mer. This allows to efficiently create an array of intersection sizes with all reference sequences on demand.

### 3.2. Post-order Lineage Tree

The core of `raxtax` is a multi-furcating tree data structure that reflects the entire lineage tree of the reference sequence set *D*. For each query, we create a new array *A* of size |*D*| to hold the normalized confidence scores from Eq. 4. The indices of *A* correspond to the leaves of the tree in post-order. Each inner node *B* of the tree also stores an integer pair (*a, b*) that contains the index interval of *A* that belongs to the rank associated with this node. After computing the confidence scores as described in Section 2 and storing them in *A*, we compute their prefix sum *A*_*p*_. To subsequently determine the clade confidence score *L*(*B*) (see Equation 5) for any clade *B* of the tree, we calculate it via *A*_*p*_[*b*] *− A*_*p*_[*a*] as can be seen Figure 2.

**Fig. 2.**
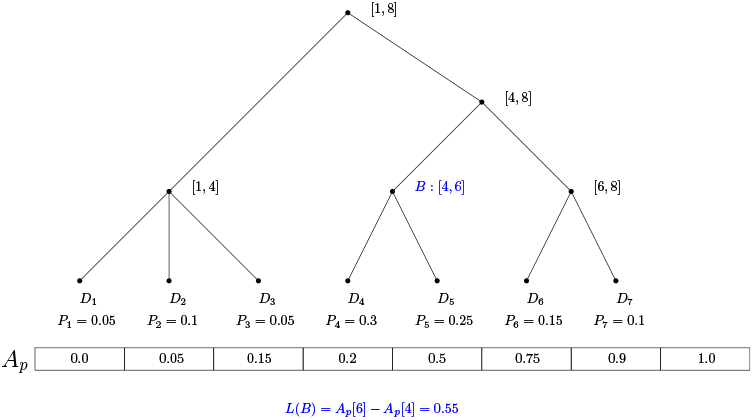
A simple lineage showing the prefix sum *A*_*p*_ and inner nodes indices. An example for node *B* is highlighted in blue.

We stop computing further *L*(*·*) values when the confidence of a node drops below a threshold of 0.005 to avoid an unnecessary evaluation of the entire tree. Thereby we only report relevant lineages.

### 3.3 The Probability of Exactly *m* Matching *k*-mers

We defined the probability mass function (PMF) *p*_*i*_(*m*) of a reference *k*-mer set *𝒦* _*i*_ having exactly *m* out of *t* matches in Equation 2. If we expand this, we obtain

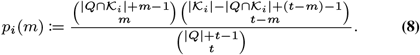

Note that for a given query, the divisor is fixed and only depends on the size of the *k*-mer set *Q* of the query sequence and *t*, that is, the number of *k*-mers to be sampled. Also note that we need to calculate the numerator for each *m ≤ t* with *m* being the only variable. By utilizing the equivalence

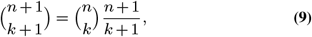

we can iteratively compute both binomial coefficients in the numerator by only using a single multiplication and division per each value of *m*.

### 3.4. Caching PMF and CMF Values. We define

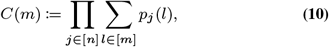

where the inner sum is the cumulative mass function (CMF) over *p*_*j*_ for a reference *k*-mer set *𝒦* _*j*_. Therefore, *C*(*m*) is the product over all CMFs for some match count *m*. Given this definition, we can compute

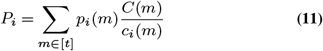

via 2*t* additional operations. Computing all PMF and CMF values has complexity *𝒪* (|*D*|*t*^2^). Using Eq. 11 decreases the additional time complexity for computing *P* from *𝒪* (|*D*|^2^*t*) to *𝒪* (|*D*|*t*). That is, the computation of best-match probabilities is reduced by a factor of |*D*|. For all but the smallest reference databases, *t ≪* |*D*|, so this caching substantially accelerates the computation.

### 3.5. Improving Runtime for Repeated Execution with the same Reference Sequences

The Lineage Tree (cf. Section 3.2) and *k*-mer-to-sequence mapping (cf. Section 3.1) are independent of any queries and can therefore be shared between runs using the same reference sequences. To this end, we save the reference database in a binary file using `bincode` (20) which conducts encoding and decoding via a tiny binary serialization strategy. This file can initially be generated and then used for further queries at a later time. Often, this saves a substantial amount of time on reference databases that comprise a large amount of sequences and/or long sequences. In our experiments with the BOLD database (21), using the binary file created by `bincode` is 2 times faster than parsing the original input.

## 4 Experimental Evaluation

We use three datasets from widely used databases: UNITE ITS (22), Greengenes 16S (23), BOLD COX1 (21, 24). In each dataset we only retained entries with complete taxonomic information and also removed duplicate sequences (Table 1). Further details about the databases can be found in Supplement 6.1. We conducted additional experiments with real-world Operational Taxonomic units (OTUs) from a large experiment of metabarcoding data from insect traps across Germany (25) and evaluated the fraction of equivalent identifications between the different tools. Among the tools, `raxtax` showed the highest agreement, with 97.66% of its classifications shared with at least one other tool. This evaluation can be found in Supplement 6.6.

**Table 1.** Databases.

| Database | UNITE | Greengenes | BOLD |
| --- | --- | --- | --- |
| Highest Taxonomic Rank | Fungi | Bacteria | Arthropoda |
| No. of sequences | 47,154 | 187,329 | 1,254,059 |
| No. of unique species | 31,479 | 629 | 136,622 |

We compare `raxtax` (v1.2.2) against four other taxonomic assignment tools: SINTAX (vsearch v2.28.1), RDP (v2.14-0), IDTAXA (DECIPHER v3.2.0) and BayesANT (v1.0).

The experiments were conducted on a a 2-socket machine with 2x Intel(R) Xeon Platinum 8260 CPUs @ 2.40GHz with 48 physical cores (96 threads) in total. Each tool was executed with 48 threads (except RDP, which can only use 2 threads) to avoid hyper-threading, unless stated otherwise.

### 4.1. Cross validation benchmarks

To evaluate `raxtax` we performed a 10-fold cross validation with random splits of the databases into 90% reference and 10% query sequences, and calculated the *F*_1_ score to assess the accuracy (TP = True Positives, MC = Missclassified, FN = False Negatives, FP = False Positives) at different taxonomic ranks.

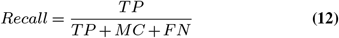

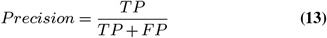

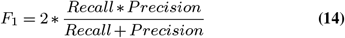

Each tool provides a confidence score for the result of each query assignment and for each taxonomic rank that ranges between 0 and 100. We evaluated our algorithm against the competing tools by setting a continuous confidence cut-off thresholds that labels all results below the respective cut-off as “not classified”. In this context, “misclassified” means that a sequence was assigned to the wrong lineage with a confidence score higher than the threshold. We then calculate the *F*_1_ score for each confidence cut-off value (Figures 3 and 4).

**Fig. 3.**
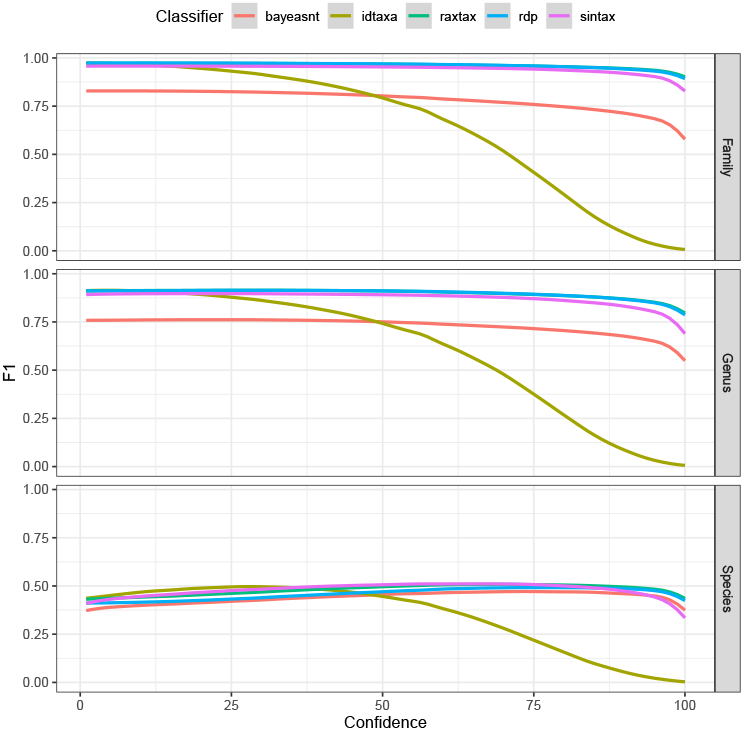
*F*_1_ scores (y-axis) for classification of UNITE sequences at the family, genus, and species level (top to bottom) where the reported confidence exceeds the confidence cut-off (x-axis).

**Fig. 4.**
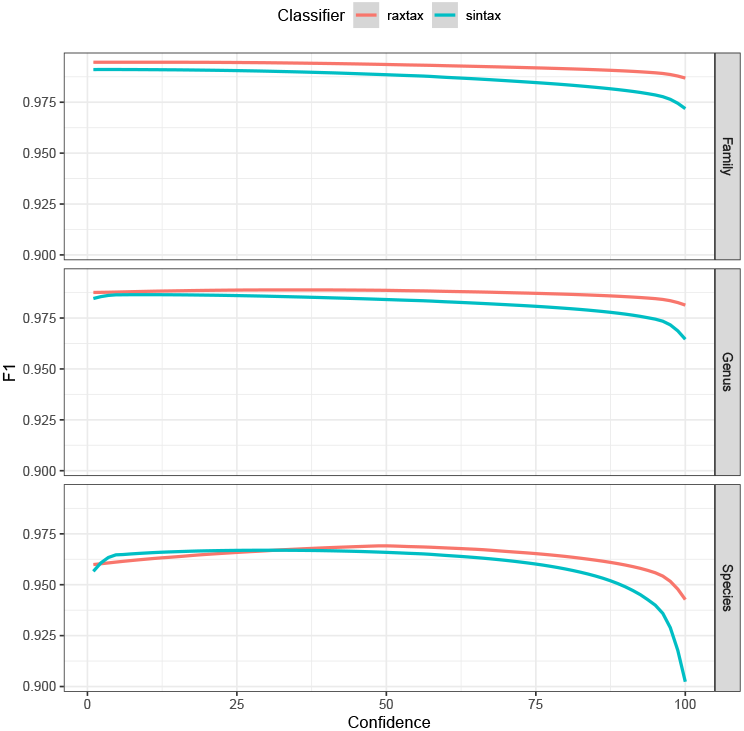
*F*_1_ scores (y-axis) for classification of BOLD sequences at the family, genus, and species level (top to bottom) where the reported confidence exceeds the confidence cut-off (x-axis).

Figure 3 shows that on the UNITE database, `raxtax`, RDP and SINTAX perform equally well at all taxonomic levels. Further, `raxtax` and RDP are indistinguishable at the family and genus level. IDTAXA was developed to circumvent over-classification. Hence, once the confidence threshold approaches values of 25-50 the computed *F*_1_ scores rapidly decline as a consequence of this conservative approach. BayesANT is only competitive at the species level.

For sequences from the BOLD database (Figure 4) only `raxtax` and SINTAX finished all 10 cross validations within the 48h time limit, so we compare only their *F*_1_ scores. Partial results including RDP and IDTAXA can be found in Supplement 6.3. The `raxtax` *F*_1_ score is consistently better at the family and genus level. At species level, the difference is statistically significant under the Wilcoxon signed-rank test with the alternative hypothesis that `raxtax` has higher *F*_1_ scores, and the matched pairs rank-biserial correlation (RBC, effect size) is large (26) (*p* = 6.3680 *×* 10^*−*77^, RBC: 0.6763). The standardized mean difference (Cohen’s d, (27)) is medium sized (*d* = 0.5549), indicating that while the *F*_1_ scores of `raxtax` are consistently higher, the differences are only marginal. In general, both tools perform exceptionally well at classifying these sequences. However, as we show in the following sections, `raxtax` is 2.7 times faster than SINTAX for the comparatively large BOLD database and exhibits growing speedups as we simultaneously increase the number of query *and* reference sequences. Results for the Greengenes database can be found in Supplement 6.2.

### 4.2. Performance Benchmarks

We measured the runtime and memory requirements of each tool for a single test (i.e., one out of the 10 cross validations) on each dataset (Table 2). We set a time limit of 48 hours –a common job time limit on clusters– to accommodate for trade-offs between accuracy and time requirements. On the BOLD dataset, only `raxtax`, SINTAX and RDP completed within the memory and time limits. We observed that RDP and BayesANT require more resources as a function of the unique species number in the reference, while SINTAX and IDTAXA performance depends on the number of query and reference sequences. Datasets will continue to grow over time, both, in terms of the species diversity they cover, and the number of query as well as reference sequences they contain. Hence, we expect that the computational resource requirements of some of the tools we tested might prohibit their future deployment.

**Table 2.**
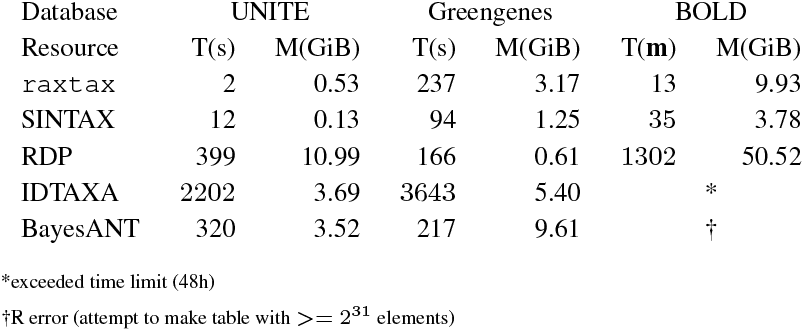
Time and memory requirements for a single cross validation. Runtimes on BOLD are in minutes instead of seconds.

### 4.3. Snapshot benchmark

In order to validate our algorithm via a more realistic setting, we used two different BOLD database snapshots that were generated eleven months apart from each other. We taxonomically classified the sequences that were added during these eleven months by treating them as query sequences and subsequently compared the inferred annotation results with the respective “true” taxonomic annotation. Given the data volume of this analysis, only `raxtax` and SINTAX were able to terminate within the 48h time limit using 48 threads. The *F*_1_ scores are shown in Figure 5. As for the 10-fold cross validation on the BOLD database (Figure 4), `raxtax` and SINTAX are equally accurate. `raxtax` again outperforms SINTAX at the family and genus level. However, the difference at species level is not statistically significant in this test. Both the effect size (Wilcoxon with two-sided alternative, *p* = 0.7635, RBC: 0.0349) and the standardized mean difference (*d* = 0.1427) are small. Here `raxtax` is 5.62 times faster than SINTAX.

**Fig. 5.**
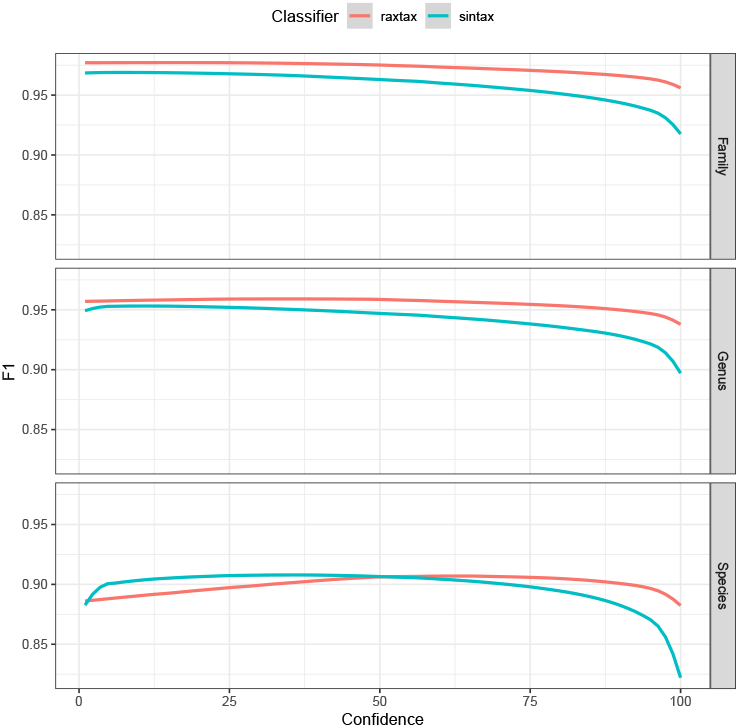
*F*_1_ scores (y-axis) for the classification of BOLD snapshots at the family, genus, and species level (top to bottom) where the reported confidence exceeds the confidence cut-off (x-axis).

### 4.4. Time and Memory Scaling

Figure 6 shows super-linear runtime scaling for both tools when we simultaneously increase the number of reference *and* query sequences. The number of threads for both tools is again fixed to 48. `raxtax` clearly scales better when we increase the number of query and reference sequences. Going from 100,000 to 1,000,000 total sequences the speedup over SINTAX increases from 1.3 to 2.9.

**Fig. 6.**
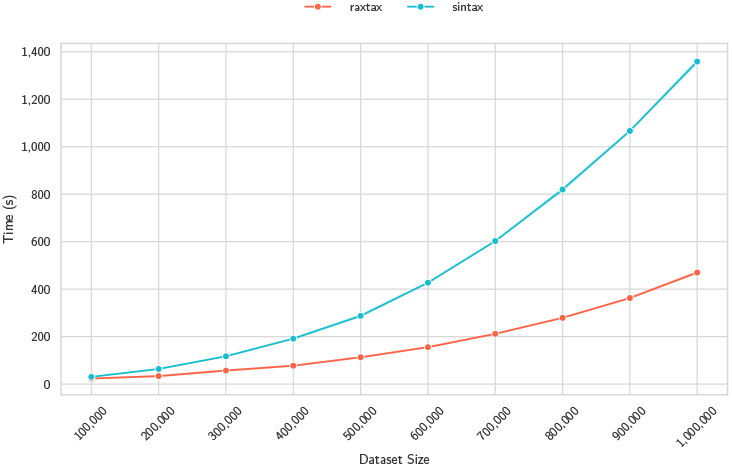
Time (y-axis) for classification of BOLD samples of different sizes (x-axis). 90% of the sample is the reference, the remaining 10% are the queries (3 random samples per sample size).

In terms of memory requirements (Figure 7), both tools exhibit a linear memory scaling as the total dataset size (no. of query and reference sequences) increases. The main memory requirements of `raxtax` (and presumably SINTAX as well) are dominated by the data structures that hold the reference database. Hence, this linear scaling is expected. SINTAX exhibits lower memory requirements and better scaling when we increase the total number of sequences. However, even for the whole BOLD database `raxtax`’s memory requirements remain below 10 GiB (see Table 2), so we argue that this is a favorable resource trade-off for using `raxtax` because of faster run times. BOLD currently contains the largest amount of barcodes for meta-barcoding projects and the memory consumption of `raxtax` increases by roughly 1GB per 150,000 sequences. Therefore, we believe that `raxtax` can be used without issues on mid-range laptops for the foreseeable future.

**Fig. 7.**
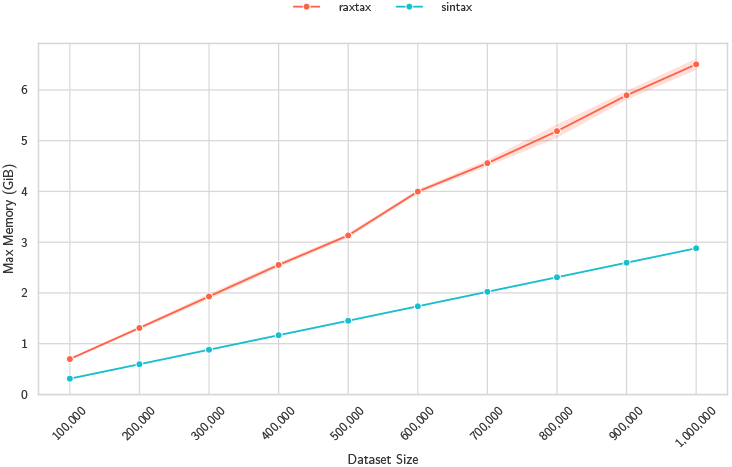
Maximum memory usage in GiB (y-axis) for classification of BOLD samples of different sizes (x-axis). 90% of the sample is the reference, the remaining 10% are the queries (3 random samples per sample size).

We also measure strong parallel efficiency for `raxtax` for a varying number of threads on samples from the BOLD database. Figure 8 shows a gradual decline in parallel efficiency from 2 threads (efficiency: 0.87) to 24 threads (efficiency: 0.74) compared to the baseline with 1 thread. Thereafter, parallel efficiency continues to rapidly deteriorate with increasing number of threads.

**Fig. 8.**
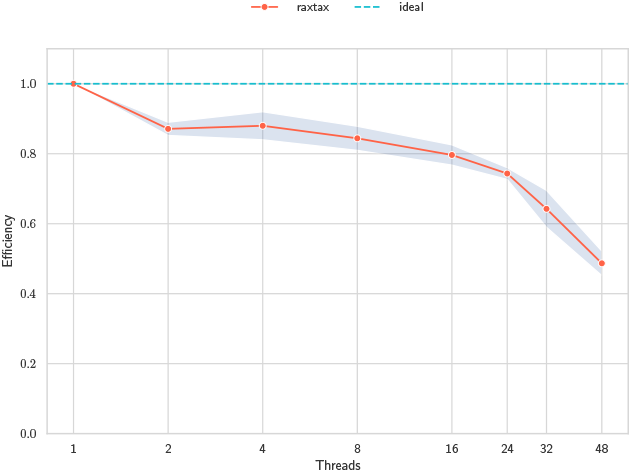
Strong self-relative efficiency and standard deviation (y-axis) over increasing thread numbers (x-axis). The reference database size is fixed at 450,000 sequences with 50,000 queries (90-10 split), and we randomly sample 5 times.

**Fig. 9.**
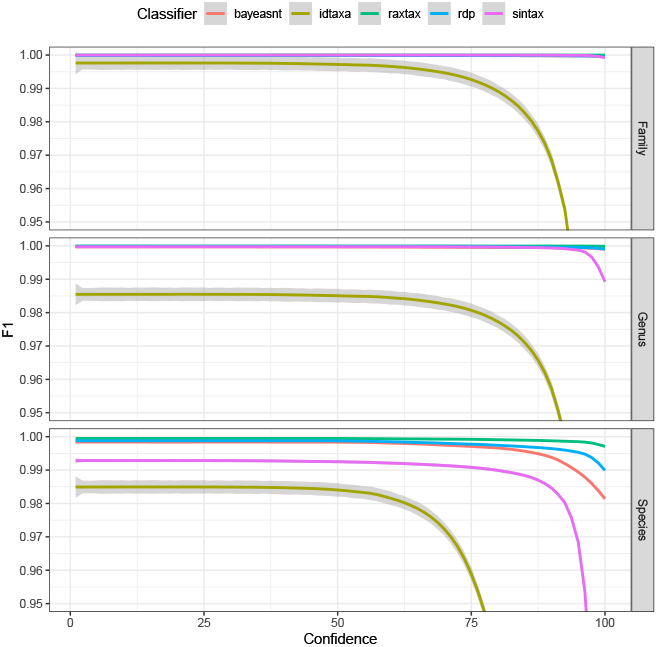
*F*_1_ scores (y-axis) for classification of Greengenes sequences at the family, genus, and species level (top to bottom) where the reported confidence exceeds the confidence cut-off (x-axis).

**Fig. 10.**
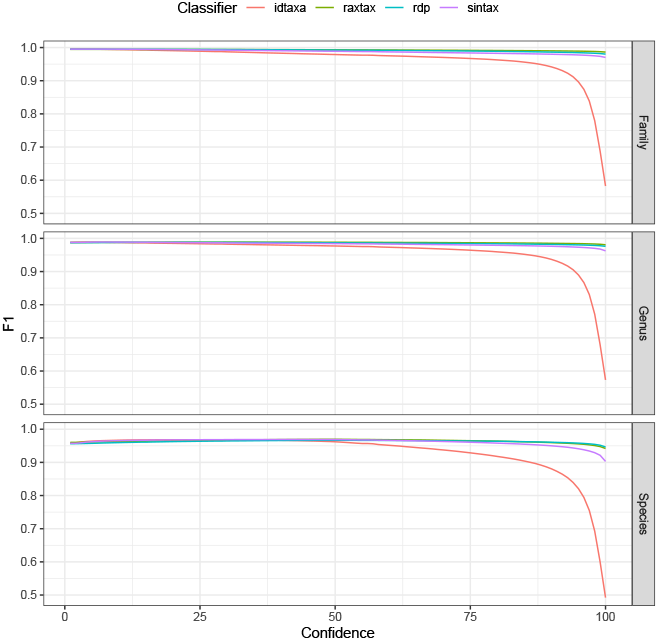
*F*_1_ scores (y-axis) for classification of BOLD sequences at the family, genus, and species level (top to bottom) where the reported confidence exceeds the confidence cut-off (x-axis).

We use explicit thread-pinning to avoid executing threads on the same physical core and schedule threads to the same socket if possible (up to 24 threads). Therefore, the more rapid decline in parallel efficiency at 32 and 48 threads is to be expected as cross-socket communication overhead is introduced. See Supplement 6.4 for more details and discussion about thread pinning and weak parallel speedup.

## 5 Conclusion

We have presented a novel analytical approach for classifying unlabeled sequences based on *k*-mer matching and derive the equations of our match scoring function for determining the best matching taxonomic lineage. Our method also introduces two additional uncertainty scores that are sensitive to an unbalanced distribution of ranks in the reference database and thereby provide users more context for drawing informed conclusions. We implemented this approach in `raxtax` as open source software. Further, we conducted a thorough code optimization to ensure that the tool is fast and efficient. An extensive evaluation of `raxtax` in conjunction with a comparison to existing tools demonstrates that we attain better or equivalent classification accuracy based on *F*_1_ scores. Further, `raxtax` can handle the ever-increasing dataset sizes in taxonomic classification and can efficiently use all available computational resources on modern hardware. We argue that the increased memory requirements compared to SINTAX are an acceptable trade-off for the reduced run-times. In the future, we aim to deploy `raxtax` as part of a comprehensive meta-barcoding pipeline for real-world queries and adapt our approach in order to apply it beyond short barcoding sequences. Finally, we intend to investigate the design of a distributed memory parallelization.

## Author contributions statement

B.B., N.W. and G.K. devised the method, N.W. implemented the code, G.K. and N.W. conceived and conducted the experiments, and analyzed the results. N.W., B.B., G.K., and A.S. wrote and reviewed the manuscript.

## ACKNOWLEDGEMENTS

We thank Robert C. Edgar for sharing his insights about developing and validating SINTAX. This work is supported by the European Union (EU) under Grant Agreement No 101087081 (Comp-Biodiv-GR).

## Data and Code Availability

This tool is available as source code and precompiled binaries at https://github.com/noahares/raxtax under a CC-NC-BY-SA license. The scripts and summary metrics used in our analyses are available at https://github.com/noahares/raxtax_paper_scripts. The source code, sequence data and summarized results of the analyses are available at https://doi.org/https://doi.org/10.5281/zenodo.15057027 (28).

## 6 Supplement

### 6.1 Detailed Database Specifications

The UNITE, Greengenes and BOLD databases are split into 90% reference sequences and 10% query sequences for all experiments. The BOLD Snapshot consists of reference sequences added up to 2023-09-29. The query sequences contain all sequences added in the following 11 months up to 2024-08-23. The number of query and reference sequences are listed in Table 3 below:

**Table 3.** Databases.

| Database | Reference Sequences | Query Sequences |
| --- | --- | --- |
| UNITE | 38,653 | 4295 |
| Greengenes | 168,596 | 18,733 |
| BOLD | 1,128,653 | 125,406 |
| BOLD Snapshot | 2,038,078 | 419,821 |

**Table 4.** Time and memory requirements for queries with real-world data.

| Tool | Time (hh:mm:ss) | Memory (GiB) |
| --- | --- | --- |
| *raxtax* | 00:04:16 | 13.39 |
| SINTAX | 01:01:49 | 6.45 |
| RDP | 09:34:31 | 113.74 |
| IDTAXA | *93:20:16 | 33.15 |
| BayesANT | † |  |

\*exceeded time limit (48h)
†R error (attempt to make table with $\geq 2^{31}$ elements)

### 6.2 Greengenes Cross Validation Benchmarks

The sequences of the Greengenes database were by far the easiest to classify. Except IDTAXA, all tools perform almost perfectly on family and genus level. Only on species level differences become apparent: While BayesANT and RDP experience slight drop-offs once the confidence cutoff approaches 100, `raxtax` maintains the highest *F*_1_-score throughout all cutoff values. SINTAX has overall lower *F*_1_-scores on species level and experiences an even stronger drop-off than BayesANT and RDP once the cutoff is beyond 90.

### 6.3 Incomplete BOLD Cross Validation Benchmarks

In Section 4.1 of the main paper we excluded RDP and IDTAXA from the results because they did not finish all 10 cross validations. Here we include them in the comparison of *F*_1_ scores for **one** of the 10 cross validation runs. As with the UNITE and Greengenes databases, RDP and `raxtax` perform equally well. IDTAXA once again cannot compete once the cutoff exceeds 50 due to its conservative approach.

### 6.4 Weak Speedup and Thread-pinning Evaluation

In Figure 11 we show the results of weak parallel speedup experiments. As `raxtax` conducts increasing work (i.e., additional queries) relative to the number of threads, we observe a gradual decrease in parallel speedup compared to the baseline with 1 thread. Overall, using thread-pinning (TP) and scheduling threads to the same socket if possible (up to 24 threads) shows better results compared to leaving the assignment of threads to the operating system. The super-linear speedup at 2 threads for `raxtax` without thread-pinning can be explained by cache coherence, as large data structures are mutually read by different threads. We attribute the gradual decline in parallel efficiency to cache contention for the L3-cache, which is shared by all physical cores of a socket. We note that the standard deviation of speedups reaches up to 0.13, indicating that the choice of reference and query sequences heavily influences the run time. In this setup, the “difficulty” of the random queries for a different number of threads can change significantly, because we add 2,000 additional queries per thread, so these results need to be interpreted with caution. Generally, we recommend using thread-pinning when running `raxtax`.

**Fig. 11.**
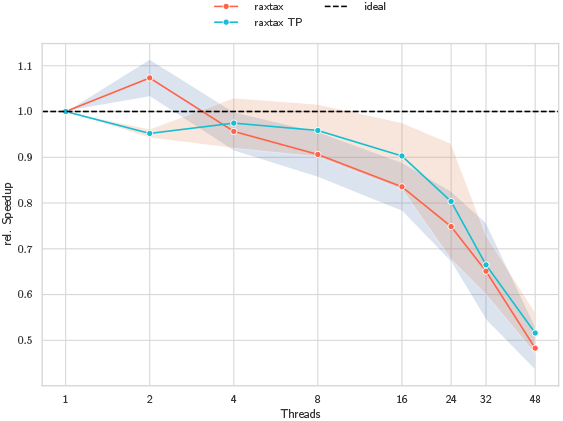
Weak self-relative speedup (y-axis) and standard deviation over increasing thread numbers (x-axis). The reference database size is fixed at 250,000 sequences, that are randomly sampled 5 times. For each additional thread a further 2,000 query sequences are included. We compare `raxtax` without thread-pinning (red) and `raxtax` with thread-pinning (cyan).

### 6.5. Using Local and Global Assignment Signals

A challenge when interpreting classification scores is that they are affected by the database composition. Thus, exclusively reporting the best match may obfuscate close runner-ups that need to also be considered. We address this challenge via our *local* assignment signal. It quantifies to which extent the assignment deviates from the expected scores based on the respective clade sizes in the given reference database. Results with a *large* local assignment signal are therefore more trustworthy. In contrast, results with a *small* local assignment signal may indicate that the confidences are inflated by an overrepresentation of that specific clade in the reference database. Note that for highly unbalanced databases, the maximum local assignment signal will be very low. We deliberately do not normalize the score to [0, 1] to increase user awareness regarding this bias.

The *global* assignment score is identical for all results computed for a query. It indicates to which degree the results can be distinguished from random matches at the sequence level. If the global assignment signal is large and roughly corresponds to the species level confidence score, the result may be interpreted as being unambiguous. In this case the query will have but a few matches with other reference sequences. If the global assignment score is small compared to the species score, there will be an increased number of matches with other reference sequences. In this case, users should inspect the results to verify or discard the assignment.

As these signals constitute a novel approach to quantify assignment confidence, we currently cannot yet provide reasonable thresholds for what should be considered as “large” or “small”. We further strongly discourage users from using them as primary scores for assignment evaluation. We recommend using large signals for justifying assignment results and small signals for critically evaluating assignment results.

### 6.6. Benchmarking tools with real-world data

We have downloaded the Operational Taxonomic units (OTUs) from a large insect metabarcoding data experiment with insect traps across Germany (25). We classified a total of 61,148 OTUs using all tools against a BOLD reference database containing 2,159,792 unique sequences. BayesANT yielded the same R error as in Table 2 of the main paper and was therefore excluded from the comparisons. Subsequently, we selected those 31,270 OTUs for which at least one of the tools inferred a confidence score of at least 0.7. We then compared these OTUs at the species level to identify areas of agreement and discrepancy among the tools and visualised the results with UpSetR ((29, 30)). All four tools agree on the species-level classification for 74.48% of the OTUs, while the remaining classifications yield different intersecting sets (Figure 12). Among the tools, `raxtax` showed the highest agreement, with at least one other tool for 97.66% of its classifications, while RDP showed the lowest agreement with 87.05%.

**Fig. 12.**
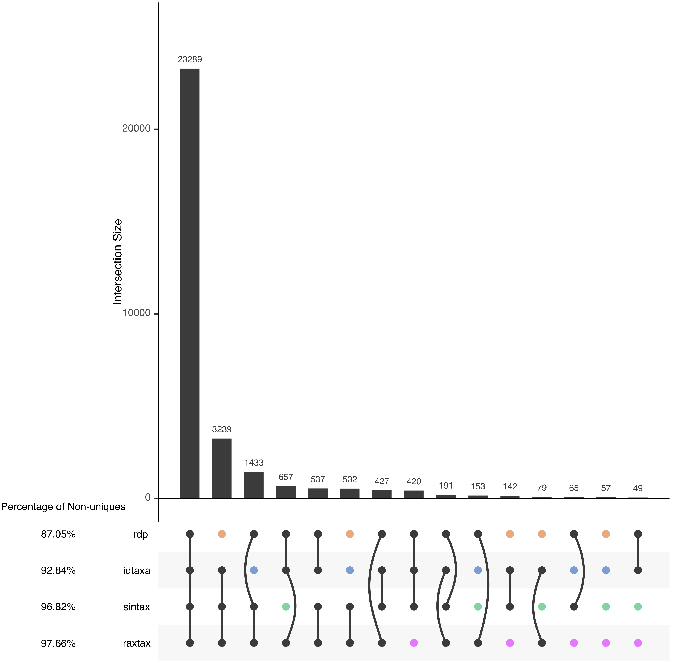
UpSet plot of shared classifications between the different tools.

Time and memory requirements are similar to the results in Table 2 of the main manuscript. `raxtax` is significantly faster than all other tools (12.75x faster than SINTAX), but requires twice as much memory as SINTAX.

